## Supplementary File 1 for "Repression of interrupted and intact rDNA by the SUMO pathway in *Drosophila melanogaster*"

**Supplementary File 1. *Drosophila melanogaster* stocks.**

| Stock | Source | Identifier |
| --- | --- | --- |
| UID rDNA unit intact | This study | N/A |
| UID rDNA unit R1 | This study | N/A |
| UID rDNA unit R2 | This study | N/A |
| UID rDNA unit R1 UID | This study | N/A |
| UID rDNA unit R2 UID | This study | N/A |
| UID rDNA unit CFP R1 | This study | N/A |
| UID rDNA unit CFP R2 | This study | N/A |
| UID Csy4 rDNA unit intact | This study | N/A |
| UASp > Csy4-Ubc9 | This study | N/A |
| UASp > Csy4-mkate2 | This study | N/A |
| UASp > Rpl135-GFP | This study | N/A |
| UASp > small hairpin white | Bloomington stock | BDSC #33623 |
| UASp > small hairpin Smt3 | M. Ninova et.al 2019 | N/A |
| UASp > small hairpin Su(var)2-10 | Bloomington stock | BDSC #32956 |
| Maternal alpha-tubulin67C > Gal4 | Bloomington stock | BDSC #7063 or #7062 |
