## Supplementary File 2 for "Repression of interrupted and intact rDNA by the SUMO pathway in *Drosophila melanogaster*"

**Supplementary File 2. Primers.**

| **For cloning of rDNA transgenes** | **Sequence (5’ to 3’)** |
| --- | --- |
| rDNA full clone 1 F | TAAGCTAGCTGTTCTACGACAGAGGG |
| rDNA full clone 1 R | GAACAATCGCCAACGTGCATTGGAG |
| rDNA full clone 2 F | TATCAAGAGTAGCCAAACACCTCG |
| rDNA full clone 2 R | CTGTCGAAGTTGTTCTCGTGAAGC |
| rDNA unit clone 1 R | TTTGAACCCGCTGTCCTCAAAGCG |
| rDNA unit clone 2 F | CGCTTTGAGGACAGCGGGTTCAAA |
| rDNA unit clone 2 R | GCGACTGAGAGAGCCATAAAAGTAGC |
| rDNA unit clone 3 F | GCTACTTTTATGGCTCTCTCAGTCGC |
| rDNA unit clone 3 R | CCATACACTGCATCTCACATTTGCC |
| rDNA unit clone 4 F | GGCAAATGTGAGATGCAGTGTATGG |
| rDNA unit clone 4 R | GTTGAGAAGTTTGCGTGTCCCCACCAT |
| rDNA unit clone 5 F | ATGGTGGGGACACGCAAACTTCTCAAC |
| rDNA unit clone 5 R | CATGACGAGAGAGTTTGCTAAGCC |
| rDNA ETS upstream R | TATAGTAGTTTTTGAACCCGCTGTCCTCAAAGCGG |
| rDNA ETS 18S F | CGGGTTCAAAAACTACTATAGGTAGGCAGTGGTTG |
| rDNA ETS 18S R | GATATTATACAATAATGATCCTTCCGCAGGTTCACCT |
| rDNA ITS1 ITS2 F | AGGTGAACCTGCGGAAGGATCATTATTG |
| rDNA ITS1 ITS2 R | GAGTTGAGGTTGTATATAACTTTATCTTGCC |
| rDNA 28S F | GGCAAGATAAAGTTATATACAACCTCAACTC |
| rDNA 28S R | CATTCGAGGGAGTAGGAATCTCGTTAATCCATTCATG |
| rDNA 28S downstream F | GATTCCTACTCCCTCGAATGGAGTGCCAGAACCCG |
| rDNA EST upstrm 2 R | GCGTATATTCCTATTATCCGCGGAGC |
| Y22 upstream EST F | GAAGCTTATCCTTTGCTTGATG |
| Y22 upstream EST R | TCCCGTGTTCAAAAAGAACTGAG |
| Y22 upstream EST with UID R | GCATCGATGTCGACTCGAGTCTCCCGTGTTCAAAAAGAACTGAG |
| Y22 EST with UID F | GACTCGAGTCGACATCGATGCCTTGGCTCCGCGGATAATAGG |
| 28S R1 R | ATGTCCACGAGCGCAACGAAAACACGTCCGGTAGGAATCTCGTTAATCCATTC |
| 28S R1 F | CGCGCATGAATGGATTAACGAGATTCCTACCGGACGTGTTTTCGTTGCGCTCG |
| R1 28S R | GTGGTTTCGCTAGATAGTAGATAGGGACAAATGTATGCTTTTCGGATCCCTC |
| R1 28S F | TAAGTTCGGAGGGATCCGAAAAGCATACATTTGTCCCTATCTACTATCTAGC |
| R1 mid R | TGGTCACTCAATTCCCACTCGTCCACTCTCCACTCATATGTGGCC |
| R1 mid F | GGCCACATATGAGTGGAGAGTGGACGAGTGGGAATTGAGTGACCA |
| 28S downstream 1 R | GAACAATATGAGAGGTCGGCAACCAC |
| 28S downstream 2 R | GGACACAATATCATATGCCTGCGTC |
| 28S R2 R | CTCTGCTCTCAAATACCCCATGATCCCCAATAAGAGAGTCATAGTTACTCC |
| 28S R2 F | TCAACGGCGGGAGTAACTATGACTCTCTTATTGGGGATCATGGGGTATTTG |
| R2 28S R | TCACTAATTAGATGACGAGGCATTTGGCTAGATCGCGGAGGTATGGAAATC |
| R2 28S F | TCTTTCGAAGATTTCCATACCTCCGCGATCTAGCCAAATGCCTCGTCATCTAA |
| R2 mid R | AGTCGACTATCGTCGCATTATCGGCGCATCCGTCGGTTGGTAAGAAGCCC |
| R2 mid F | GGGCTTCTTACCAACCGACGGATGCGCCGATAATGCGACGATAGTCGACT |
| R1 F | CGGACGTGTTTTCGTTGCGCTCG |
| R1 R | ATGTATGCTTTTCGGATCCCTCCG |
| R2 F | TTGGGGATCATGGGGTATTTGAGAG |
| R2 R | GATCGCGGAGGTATGGAAATCTTC |
| 28S eCFP R1p R | GGTGAACAGCTCCTCGCCCTTGCTCACCATGTAGGAATCTCGTTAATCCATTC |

| 28S eCFP R1p F | CGCGCATGAATGGATTAACGAGATTCCTACATGGTGAGCAAGGGCGAGGAGC |
| --- | --- |
| eCFP 28S R1p R | TGTGGTTTCGCTAGATAGTAGATAGGGACAATTACTTGTACAGCTCGTCCATG |
| eCFP 28S R1p F | CACTCTCGGCATGGACGAGCTGTACAAGTAATTGTCCCTATCTACTATCTAGC |
| 28S eCFP R2p R | CGGTGAACAGCTCCTCGCCCTTGCTCACCATTAAGAGAGTCATAGTTACTCC |
| 28S eCFP R2p F | GTCAACGGCGGGAGTAACTATGACTCTCTTAATGGTGAGCAAGGGCGAGGAGC |
| eCFP 28S R2p R | GTCACTAATTAGATGACGAGGCATTTGGCTATTACTTGTACAGCTCGTCCATG |
| eCFP 28S R2p F | ACTCTCGGCATGGACGAGCTGTACAAGTAATAGCCAAATGCCTCGTCATCTAA |
| 28S R1 adaptor R | CCGCGCTCGAGCCCTATAGTGAGTCGTATTAGGTAGGAATCTCGTTAATCCATTC |
| 28S R1 adaptor F | TACCTAATACGACTCACTATAGGGCTCGAGCGCGGACGTGTTTTCGTTGCGCTCG |
| R1 28S adaptor R | GACAAGAAGCATGGATCCGGCGCGAACTCGAGATATGTATGCTTTTCGGATCCCTC |
| R1 28S adaptor F | ACATATCTCGAGTTCGCGCCGGATCCATGCTTCTTGTCCCTATCTACTATCTAGC |
| 28S R2 adaptor R | CCCCAACGCTCGAGCCCTATAGTGAGTCGTATTAGTAAGAGAGTCATAGTTACTCC |
| 28S R2 adaptor F | CTCTTACTAATACGACTCACTATAGGGCTCGAGCGTTGGGGATCATGGGGTATTTG |
| R2 28S adaptor R | GGCTAGAAGCATGGATCCGGCGCGAACTCGAGATGATCGCGGAGGTATGGAAATC |
| R2 28S adaptor F | GCGATCATCTCGAGTTCGCGCCGGATCCATGCTTCTAGCCAAATGCCTCGTCATCTAA |
| P36 rDNA bgning GA F | GGGTTTTATTAACTTACATACATACTAGAATTGGGCCGAAGCTTATCCTTTGCTTGATG |
| rDNA end P36 GA R | GTTTTATTTTTAATAATTTGCGAGTACGCAAAGCTTGGCTGCATGGCGCGGGACACAATATCATATGCCTGCGTC |
| #557 EXT. R | AATCGTTAATTTCTCATACTAGAATATTGACGCTCCATACACTGCATCTCACATTTGCC |
| Csy4 rDNA F | GTTTCTCAGTTCTTTTTGAACACGGGAGTTCACTGCCGTATAGGCAGGACT |
| Csy4 rDNA R | GAGCCAAGGCATCGATGTCGACTCGAGTCCTGCCTATACGGCAGTGAACTCCC |
| **Primers for qPCR** | **Sequence (5’ to 3’)** |
| UID qPCR F | GACTCGAGTCGACATCGATGC |
| UID qPCR R | TTGCCTGCCACCAAAAATTAAC |
| rRNA-R1 qPCR F | CGGACGTGTTTTCGTTGCGCT |
| rRNA-R1 qPCR R | CCTTAGCGGTGACTACCACCAATAA |
| R1 qPCR F | AGATCGGAGTCAATGTTTGCTCT |
| R1 qPCR R | GAGCCAAAACTCCCTTTAACGAC |
| rRNA-R2 qPCR F | TTGGGGATCATGGGGTATTTGA |
| rRNA-R2 qPCR R | TGCTTGTAGTTCCAATATGAATAAATTTCC |
| R2 qPCR F | GGTCCTTTAACAGCAAGAGAGGA |
| R2 qPCR R | CTCCTTATAATCGCCCCTCTGTC |
| ETS qPCR F | ATTACCTGCCTGTAAAGTTGG |
| ETS qPCR R | CCGAGCGCACATGATAATTCTTCC |
| 18S qPCR F | CTTACCAGGTCCGAACATAAG |
| 18S qPCR R | GACAAATCACTCCACGAACTA |
| Kalahari qPCR F | TGGTCTACGACATGGGCAAC |
| Kalahari qPCR R | TGGGAATCGTCAGGGGTATC |
| rp49 qPCR F | CCGCTTCAAGGGACAGTATCTG |
| rp49 qPCR R | ATCTCGCCGCAGTAAACGC |
| H3K9me3 enriched region F | GTTGCTGGTAGCACGACGTA |
| H3K9me3 enriched region R | GTCCTTTTCGTCCAGTCTGC |
| Smt3 qPCR | TAACGAGGTAGGTCCAGCAC |
| Smt3 qPCR | CTTTTGCGGAGCGTCGAATG |
| dgrn qPCR F | TGAATCGCTACTGCGAGGAAG |
| dgrn qPCR R | TTCTGCGGCGAATTCTACGAG |
| Ulp1 qPCR F | GCCGATAAGACTGCGATTTC |
| Ulp1 qPCR R | AGATCGATCAGGGCATCAAC |
| velo qPCR F | ACTGCCAGTGATCGAGTTCC |
| velo qPCR R | CCCTCCTCCTCCTCAAACTC |
| Qjt qPCR F | TGCGGAAGTCAATGATCAAG |
| Qjt qPCR R | TGGACCAGGGATCGTAGAAC |
| Topors qPCR F | AGCACGACATAGCCAACTCC |
| Topors qPCR R | CATGGAATTCTCTGCCGAAT |
| Nse2like qPCR F | CAAAATGGACCCGAAATCC |
| Nse2like qPCR R | TAGCAACTTCTCCCCGTCAT |
| CG13773 qPCR F | CAAGAAAGAGGACCCAGATGA |
| CG13773 qPCR R | CCATTCTTGCGTTTTGTTGA |
| l(2)37Cg qPCR F | CCACCAAAGCACAACAGATG |
| l(2)37Cg qPCR R | CTTCTCGGCATCCTGCTAAC |
| CG3756 qPCR F | CAGCTAAACGACGCTGTGAC |
| CG3756 qPCR R | AAGATGACATCGGGCTTCAG |
| Udd qPCR F | AACGACGAAGTTAGGCAAGAA |
| Udd qPCR R | TAGTGGGATGGACGCAGTTT |
| Nop60B qPCR F | ATTCAGTCTGAGCGCGATG |
| Nop60B qPCR R | ACTCGTCCTTGTGGTTTTCG |
| CG4038 qPCR F | ACGAGGTGGAGGCGGATT |
| CG4038 qPCR R | CGCCTCCTCGTCCTCTGTT |
| NHP2 qPCR F | AAGTAGAGCGCTCCGAGGAT |
| NHP2 qPCR R | ATGGCGTTCACAAAGATCAG |
| Fib qPCR F | CAAGACCGTCACCATCGAG |
| Fib qPCR R | TCGGATCCAGGTACAAAGTTC |
| CG1785 qPCR F | TGAGGAGGAGGACCTGAATG |
| CG1785 qPCR R | CTAGGTCTGGCTCCTCGAAC |
| mbm qPCR F | CAGCAAATGACTGCCTTTCA |
| mbm qPCR R | TTGTCCTTTTTGGGTTTTGC |
| Dp1 qPCR F | AAGTTCCCTGATCGTGATGC |
| Dp1 qPCR R | CTTCATTCTCGCCTCCATTC |
| Hrb27C qPCR F | AACAATACGGCTACGGCAAC |
| Hrb27C qPCR R | GCTGAGGTCCAGCGTAGTTC |
| Iswi qPCR F | GATCGTGGAGAGAGCAGAGG |
| Iswi qPCR R | AACTGATTGGAGCGGTTGTC |
| apt qPCR F | TGTTCATGGAGATGCAGAGC |
| apt qPCR R | ACTAGGTCGCGACTGTGCTT |
| Nlp qPCR F | CCGACGTGGAGTTCTACGAG |
| Nlp qPCR R | CATCGTCCTTGATGTTGTGC |
| mod qPCR F | ACGTCAACTGGAGCAGAAGC |
| mod qPCR R | ATTTCTCGGTTGGGACACAG |
| sle qPCR F | ACTCCCAAAGTGGCAGTCAG |
| sle qPCR R | CCGGTACAATCAATGGCTTC |
| brat qPCR F | CTTCGAATGCCAGAGCTACC |
| brat qPCR R | CGCCTCAAAACCCTTAATCA |
| Top1 qPCR F | ACAACGAGGTTGAGGTGGAG |
| Top1 qPCR R | CGTGTTGAGACGATCGAAGA |
| **DNA targets for HCR FISH** | **Sequence (5’ to 3’)** |
| UID | GTTTCTCAGTTCTTTTTGAACACGGGAGACTCGAGTCGACATCGATGCCTTG |
| ETS | TTTTTGGTGGCAGGCAAATATTAGTTTATTACCTGCCTGTAAAGTTGGATTA |
| ETS | AAAATTTATATATAAATTTGGAAGAATTATCATGTGCGCTCGGTTTTATGTT |
| Vasa mRNA | AGAATCATGACGCATGTAACTATGCGTCCAGAACATCAGACATTGATGTTTT |
| Vasa mRNA | CTAGATCGGGAGGAACGCGGCGGTGAACGTCGTGGAAGACTAGATCGGGAGG |
| Vasa mRNA | CGAGAGTTTTATATTCCTCCCGAGCCGTCCAACGATGCAATTGAGATATTCA |
| Vasa mRNA | TCCCCAACAAGGGAGCTGGCAATTCAGATCTTTAACGAGGCGAGGAAGTTTG |
| Vasa mRNA | CTTCTGGATTTCGTGGATCGGACCTTTATCACGTTTGAAGACACTCGATTCG |
| Vasa mRNA | TTAGCTTCCTTCCTGTCAGAAAAGGAGTTTCCGACGACCTCCATTCATGGCG |
| Vasa mRNA | CGTCTCCAGAGTCAACGCGAGCAGGCCTTGCGTGATTTCAAGAACGGCTCTA |
| Vasa mRNA | ATTGGACGTACAGGTCGTGTAGGCAATAATGGACGAGCCACAAGCTTCTTTG |
| Vasa mRNA | AGTGGCATAGCTTCCGGGATTCATTTTTCGAAATACAACAACATACCGGTTA |
| Vasa mRNA | TTCGAGTCATATCTAAAGATCGGTATTGTTTACGGAGGCACCTCGTTCAGAC |
| Vasa mRNA | CAAAACGAGTGCATTACCAGAGGCTGCCATGTAGTGATCGCCACTCCGGGAC |
| Vasa mRNA | GCCACGTTTCCAGAAGAAATCCAAAGAATGGCCGGCGAATTCCTTAAAAATT |
| Vasa mRNA | ATATACGAAGTTAATAAGTACGCCAAGCGATCCAAGCTAATAGAAATCCTTT |
| Vasa mRNA | GAGCAAGCAGATGGCACCATTGTGTTTGTGGAGACAAAGCGTGGCGCCGACT |
| Vasa mRNA | CCTGAAAAGGATCGAGCTATTGCTGCGGACTTGGTAAAAATCTTGGAGGGAT |
| Vasa mRNA | GGCCAGACTGTTCCGGACTTTCTACGCACCTGTGGTGCCGGCGGTGATGGGG |
| Vasa mRNA | TACTCCAATCAAAATTTCGGTGGGGTGGATGTGCGTGGCAGGGGAAATTACG |
| Vasa mRNA | ATTCAAAAGTGTTCCATACCAGTAATATCCTCTGGTCGAGACCTAATGGCGT |
| Vasa mRNA | TTGTTGGAGGATCCCCATGAACTGGAGCTTGGAAGACCCCAGGTAGTGATTG |
| Vasa mRNA | GTTTTCGTCGCCATTGGCATTGTAGGCGGAGCTTGCTCTGATGTGAAGCAGA |
| **Primers for RNAi** | **Sequence (5’ to 3’)** |
| T7 Smt3 RNAi 1 F | TAATACGACTCACTATAGGGCAGCTTCAACAAGCAACCAA |
| T7 Smt3 RNAi-1 R | TAATACGACTCACTATAGGGTAATCTTATGGAGCGCCACC |
| T7 Smt3 RNAi 2 F | TAATACGACTCACTATAGGGACGCACACAGACGCATTTAG |
| T7 Smt3 RNAi 2 R | TAATACGACTCACTATAGGGAGGAAGCTGATGAACGCCTA |
| T7 dUbc9 RNAi F | TAATACGACTCACTATAGGGAGACCACGTCCGGCATTGCTATTACAC |
| T7 dUbc9 RNAi R | TAATACGACTCACTATAGGGAGACCACGGCCTGGGCACGCACGCGC |
| T7 Su(var)2-10 RNAi F | TAATACGACTCACTATAGGGCAGGAAGTCTACGCCCAGTC |
| T7 Su(var)2-10 RNAi R | TAATACGACTCACTATAGGGGCGTGAAGTAGAAAGGCACCTC |
| T7 GFP RNAi F | TAATACGACTCACTATAGGGACGTAAACGGCCACAAGTTC |
| T7 GFP RNAi R | TAATACGACTCACTATAGGGTGTTCTGCTGGTAGTGGTCG |
| T7 dgrn dsRNA F | CTAATACGACTCACTATAGGGGAGTCCAGTAGAAGTGATAGA |
| T7 dgrn dsRNA R | CTAATACGACTCACTATAGGGAAGTAAATGCGAAAGAATTGAC |
| T7 Ulp1 dsRNA F | CTAATACGACTCACTATAGGGTTGGAAATATCCCAGGAAAACA |
| T7 Ulp1 dsRNA R | CTAATACGACTCACTATAGGGATTTCTAAGCCCATCGAAATG |
| T7 velo dsRNA F | CTAATACGACTCACTATAGGGGAGGACTTCGTCTGCCTCAC |
| T7 velo dsRNA R | CTAATACGACTCACTATAGGGCGGGGTTATGGTGGTGTTAC |
| T7 qjt dsRNA F | CTAATACGACTCACTATAGGGCTGATTCCGCACTTTCCACT |
| T7 qjt dsRNA R | CTAATACGACTCACTATAGGGTGGCGCTAGAGTTTTTGACC |
| T7 Topors dsRNA F | CTAATACGACTCACTATAGGGCAATCCTGGTGCCACACAT |
| T7 Topors dsRNA R | CTAATACGACTCACTATAGGGTTGTGCTGGGCCCCATAA |
| T7 CG42299, CG42300 dsRNA F | CTAATACGACTCACTATAGGGGTCCGAGGAAGTCTGCAA |
| T7 CG42299, CG42300 dsRNA R | CTAATACGACTCACTATAGGGCGCACCGGGCATAGGA |
| T7 CG13773 dsRNA F | CTAATACGACTCACTATAGGGCTCGCAATGGCCAAGATACT |
| T7 CG13773 dsRNA R | CTAATACGACTCACTATAGGGGAGGTGTTGAAGACCCGGTA |
| T7 l(2)37Cg dsRNA F | CTAATACGACTCACTATAGGGCTGGCTGGTGATGAGAAAAA |
| T7 l(2)37Cg dsRNA R | CTAATACGACTCACTATAGGGCCACTTTCATTGCATTAAACTC |
| T7 CG3756 dsRNA F | CTAATACGACTCACTATAGGGATCGAAAAGGTGTACATATACAA |
| T7 CG3756 dsRNA R | CTAATACGACTCACTATAGGGTGAGAAACAGTTCTGTAATAGG |
| T7 udd dsRNA F | CTAATACGACTCACTATAGGGGCCATCGCCTTTGAAAACT |
| T7 udd dsRNA R | CTAATACGACTCACTATAGGGGATAGAATAGCATTTAATGAATCG |
| T7 Nop60B dsRNA F | CTAATACGACTCACTATAGGGTGCTATGGTGCCAAGATTAC |
| T7 Nop60B dsRNA R | CTAATACGACTCACTATAGGGGGGTTCACTGGAGCCATTT |
| T7 CG4038 dsRNA F | CTAATACGACTCACTATAGGGATCCCCTTGGGCAACTATGT |
| T7 CG4038 dsRNA R | CTAATACGACTCACTATAGGGTTTGTGAAGGCTTTCTTGGC |
| T7 NHP2 dsRNA F | CTAATACGACTCACTATAGGGGAAACCGATGGCAGGTAAAA |
| T7 NHP2 dsRNA R | CTAATACGACTCACTATAGGGGCAGACAGCTCCTCCTTGAC |
| T7 CG7637 dsRNA F | CTAATACGACTCACTATAGGGTGATGTACACAATTAACGAAAAC |
| T7 CG7637 dsRNA R | CTAATACGACTCACTATAGGGTAAATGGGCTCCGGCTTC |
| T7 Fib dsRNA F | CTAATACGACTCACTATAGGGCAATGGCGAGAAGATTGAGT |
| T7 Fib dsRNA R | CTAATACGACTCACTATAGGGAGCGTGAGCTGCTCTTG |
| T7 CG1785 dsRNA F | CTAATACGACTCACTATAGGGCACTGCACACAGCTGGAAAT |
| T7 CG1785 dsRNA R | CTAATACGACTCACTATAGGGTTTAGGCGACGACGATCTCT |
| T7 mbm dsRNA F | CTAATACGACTCACTATAGGGGTTACGCCAGAAGAACCTAC |
| T7 mbm dsRNA R | CTAATACGACTCACTATAGGGCATATGCCACACACATCCAC |
| T7 Dp1 dsRNA F | CTAATACGACTCACTATAGGGAAGATTCGCGAGGTGAAG |
| T7 Dp1 dsRNA R | CTAATACGACTCACTATAGGGTATCGCTCTTGGAGTCGC |
| T7 Hrb27C dsRNA F | CTAATACGACTCACTATAGGGCTTCGGGGACATCATTGACT |
| T7 Hrb27C dsRNA R | CTAATACGACTCACTATAGGGGCGGGACTTCTTCTTCTCCT |
| T7 Iswi dsRNA F | CTAATACGACTCACTATAGGGTCAGAGCCGTCTGCCTTATT |
| T7 Iswi dsRNA R | CTAATACGACTCACTATAGGGAATTAAACACATCGGGCAGC |
| T7 apt dsRNA F | CTAATACGACTCACTATAGGGTTCTGGCCTAACCAGCACTT |
| T7 apt dsRNA R | CTAATACGACTCACTATAGGGACGACAACTGAAGCCGAGTT |
| T7 Nlp dsRNA F | CTAATACGACTCACTATAGGGGAAGTAAACTGCTGACCGCC |
| T7 Nlp dsRNA R | CTAATACGACTCACTATAGGGAGGAAGTCCGGCTTAGCTTT |
| T7 mod dsRNA F | CTAATACGACTCACTATAGGGTCAAGGACGATGAGGGTTTC |
| T7 mod dsRNA R | CTAATACGACTCACTATAGGGGCTGTGGCCGTATTTATGGT |
| T7 sle dsRNA F | CTAATACGACTCACTATAGGGGTAGAAACTCAGCAACTGATT |
| T7 sle dsRNA R | CTAATACGACTCACTATAGGGTCTCCGGCTTGGGGGTT |
| T7 brat dsRNA F | CTAATACGACTCACTATAGGGAAGACCGAATTGCTCAAGG |
| T7 brat dsRNA R | CTAATACGACTCACTATAGGGCGCCTGCAGCTGTTGA |
| T7 Naa40 dsRNA F | CTAATACGACTCACTATAGGGAAGAATCCCCTCGAATCTCT |
| T7 Naa40 dsRNA R | CTAATACGACTCACTATAGGGGATCTCGTCCTTGACGTAG |
| T7 Top1 dsRNA F | CTAATACGACTCACTATAGGGGGAGGAGGAGAAGCGTG |
| T7 Top1 dsRNA R | CTAATACGACTCACTATAGGGGCGCCGCTTGATCATG |
