## Supplementary File 4 for "Repression of interrupted and intact rDNA by the SUMO pathway in *Drosophila melanogaster*"

| **Gene name** | **Gene symbol** | **Protein function** | **R1 up-regulation** | **R2 up-regulation** | **KD efficiency** |
| --- | --- | --- | --- | --- | --- |
| **SUMOylation pathway proteins** | | | | | |
| *smt3* | CG4494 | Drosophila SUMO protein | 6802,9^***^ | 1432,4^***^ | 10,5^***^ |
| *dgrn* | CG10981 | SUMO-Targeted ubiquitin ligase; recognizes SUMOylated substrates and catalyze their ubiquitylation | 0,9 | 1,1 | 11,9^***^ |
| *Ulp1* | CG12359 | SUMO-specific isopeptidase; catalyzes both SUMO maturation and SUMO deconjugation | 7,7^***^ | 25,5^***^ | 11,0^***^ |
| *velo* | CG10107 | SUMO-specific protease; predicted to have SUMO-specific endopeptidase and isopeptidase activities | 1,2 | 3,5^***^ | 5,3^**^ |
| *qjt* | CG13732 | SUMO E3 ligase; involved in heterochromatin DNA repair of DSB, subunit of the Smc5/6 complex | 1,1 | 0,9 | 1,3 |
| *Topors* | CG15104 | SUMO E3 ligase; RING-finger protein, putative dual Ubiquitin/SUMO ligase | 1,1 | 1,3^*^ | 2,1^***^ |
|  | CG42299  CG42300 | Predicted to have SUMO E3 ligase activity; predicted components of the SMC5/6 protein complex | 0,9 | 1,3 | 7,3* |
| **Histone methyltransferases (HMTs)** | | | | | |
| *Egg*  *(SetDB1)* | CG12196 | Histone H3-K9 methylase; colocalizes with chromocenter | 1,0 | 0,7^**^ | 7,1^***^ |
| *Su(var)*  *3-9* | CG43664 | Histone H3-K9 methylase; colocalizes with heterochromatin and polytene chromosome chromocenter; known as nucleolar organization regulator | 2,4^***^ | 1.9^**^ | 5,4^***^ |
| *G9a* | CG2995 | Histone H3-K9 methylase; colocalizes with euchromatin and polytene chromosome interbands; orthologous to human EHMT1 EHMT2 | 1,1 | 0.9 | 2,7^**^ |

| **Gene name** | **Gene symbol** | **Encoded protein information** | | | | **R1 up-regulation** | **R2 up-regulation** | **KD efficiency** |
| --- | --- | --- | --- | --- | --- | --- | --- | --- |
|  |  | **Protein function/nucleolus localization** | **SUMOylation reported** | **Association with GO term ‘nucleolus’** | **Presence of SUMOylation motif(s)** |  |  |  |
| **Proteins involved in rRNA transcription** | | | | | | | | |
|  | CG13773 | Predicted to contribute to RNA polymerase I activity, orthologous to human RPA43 | MS (Ninova et al., unpublished) | + | + | 8,6^***^ | 10,0^***^ | 13,6^***^ |
|  | CG3756 | Predicted to contribute to RNA polymerase I and III activity, orthologous to human POLR1C | IP/Western blot, MS (Ninova et al., unpublished) | + | + | 6,8** | 32,2** | 6,2^**^ |
| *udd* | CG18316 | Member of SL1-like complex; involved in positive regulation of RNA polymerase I transcription; localizes in nucleolus | IP/Western blot, MS (Ninova et al., unpublished) | + | + | 1,6^**^ | 1,2^*^ | 5,2^***^ |
| **Proteins involved in rRNA processing and ribosome biogenesis** | | | | | | | | |
| *Nop60B* | CG3333 | Predicted H/ACA small nucleolar ribonucleoproteins (snoRNPs);  localizes to the nucleolus | MS (Handu et al., 2015, Ninova et al., unpublished) | + | + | 1,5^*^ | 0,9 | 9,1^***^ |
|  | CG4038 | Predicted H/ACA small nucleolar ribonucleoproteins (snoRNPs);  localizes to the nucleolus | MS (Handu et al., 2015, Ninova et al.,  unpublished) | + | + | 2,0^***^ | 0,8 | 23,1^***^ |
| *NHP2* | CG5258 | Predicted H/ACA small nucleolar ribonucleoproteins (snoRNPs) | MS (Handu et al., 2015) | + | + | 2,3^**^ | 1,0 | 18,7^***^ |
|  | CG7637 | Predicted H/ACA small nucleolar ribonucleoproteins (snoRNPs) | MS (Ninova et al., unpublished) | + | - | 1,3 | 0,7^*^ | N/A |
| *Fib* | CG9888 | Predicted rRNA 2'-O-methyltransferase activity; localizes to the nucleolus | MS (Ninova et al., unpublished) | + | + | 4,4^***^ | 3,1^***^ | 21,5^***^ |
|  | CG1785 | Predicted to be involved in rRNA processing and ribosome biogenesis; orthologous to human NOP53 | MS (Ninova et al., unpublished) | + | + | 0,9 | 1,6 | 7,1^***^ |
| *mbm* | CG11604 | Zinc finger protein; involved in ribosome biogenesis; localizes to the nucleolus | MS (Ninova et al., unpublished) | + | + | 2,6** | 20,5*** | 9,1^***^ |
| **Proteins localized to the nucleolus/binding rDNA** | | | | | | | | |
| *Dp1* | CG5170 | Single-stranded nucleic acid binding protein;  binds ribosomal IGS *in vitro* (Watase and Yamashita, 2018) | MS (Ninova et al., unpublished) | - | - | 1,1 | 0,8* | 11,5^***^ |
| *Hrb27C* | CG10377 | Single-stranded nucleic acid binding protein;  binds ribosomal IGS *in vitro* (Watase and Yamashita, 2018) | MS (Ninova et al., unpublished) | - | + | 1,4 | 1,3** | 14,8^***^ |
| *Iswi* | CG8625 | Chromatin remodeling factor, component of ToRC complex; binds ribosomal IGS *in vitro* (Watase and Yamashita, 2018) | MS (Handu et al., 2015) | - | + | 1,4* | 1,1 | 12,5^***^ |
| *Nlp* | CG7917 | Chromatin remodeling factor;  localizes to the nucleolus | MS (Ninova et al., unpublished) | + | + | 1,0 | 1,0 | 32,7^***^ |
| *mod* | CG2050 | DNA-binding protein;  localizes to the nucleolus | MS (Ninova et al., unpublished) | + | + | 1,4 | 1,1 | 18,9^***^ |
| *sle* | CG12819 | Involved in nucleolus organization;  localizes to the nucleolus | MS (Handu et al., 2015, Ninova et al.,  unpublished) | + | + | 0,4 | 0,8 | 15,7^***^ |
| ***Proteins involved in rRNA transcription regulation*** | | | | | | | | |
| *brat* | CG10719 | mRNA binding protein;  involved in rRNA transcription inhibition | MS (Ninova et al., unpublished) | - | + | 4,2 | 1,3 | 2,9^*^ |
| *Naa40* | CG7593 | Predicted to have acetyltransferase activity and to be involved in regulation of chromatin silencing at rDNA | MS (Handu et al., 2015, Ninova et al.,  unpublished) | - | + | 0,6 | 1,6 | N/A |
| Top1 | CG6146 | DNA topoisomerase type I; localizes to the nucleolus; involved in rDNA transcription regulation in yeast | MS (Ninova et al., unpublished) | + | + | 2,1* | 1,2 | 6,3^*^ |
